# Task-related modulation of event-related potentials does not reflect changes to sensory representations

**DOI:** 10.1101/2024.01.20.576485

**Authors:** Reuben Rideaux

## Abstract

Attention supports efficient perception by increasing the neural signals of targets while supressing those of distractors. Decades of work studying the event-related potentials of electroencephalography (EEG) recordings have established our understanding of attention in the human brain, but many aspects of this phenomenon remain unknown. Several recent studies suggest that multivariate analyses may provide new insights into how attention shapes the neural representations of stimuli; however, it is unclear whether the increased multivariate decoding accuracy associated with task relevance represents a change in the stimulus representation or an additional cognitive process. To understand what the change in multivariate information that is associated with task relevance reflects, here we used inverted encoding to characterize how task relevance shapes the neural representation of space and colour. For both spatial and feature-based tasks, we found that the change in the EEG recordings associated with task relevance is not related to the stimulus representation. Rather, our findings indicate that this phenomenon reflects an additional cognitive process, such as target recognition.

## INTRODUCTION

To efficiently perceive our environment, the brain selectively filters sensory information using a variety of mechanisms, collectively referred to as *attention*. Attention can be captured by salient stimulus properties (i.e., *bottom-up attention*) or directed by goal-driven behaviour (i.e., *top-down attention*). Top-down attention is supported by a network of regions across frontal and parietal cortices (Buschman & Miller, 2007; Suzuki & Gottlieb, 2013; Ungerleider, 2000; Womelsdorf et al., 2008), which influence processing in sensory cortices and thalamus (Brefczynski & DeYoe, 1999; Gandhi et al., 1999; Luck et al., 1997; Motter, 1993; Treue & Maunsell, 1996), for example, by adjusting the tuning profiles of neurons to increase sensitivity for attended features (David et al., 2008; Womelsdorf et al., 2008). This increases the signal-to-noise ratio of neural activity associated with attended, relative to unattended, features (Desimone & Duncan, 1995), and is associated with increased test-retest reliability (Arazi et al., 2019).

Top-down attention is typically categorized into two forms: spatial (attending to a particular spatial location) and feature-based (attending to a specific sensory feature, such as the colour red); however, other attentional processes have been identified, such as object-based attention (Chen, 2012; Duncan, 1984). Note, here we refer to covert spatial attention, i.e., fixating one location while attending to another. While spatial and feature-based attention appear to have common phenomenology, such as increasing the gain of neurons tuned to the attended location/feature (Maunsell & Treue, 2006; Treue & Maunsell, 1996), theoretical modelling work and empirical psychophysical evidence in humans suggests that they are supported by distinct neural mechanisms (Herrmann et al., 2012; Ling et al., 2009).

Understanding the neural mechanisms of attention remains a central pursuit in cognitive neuroscience. Among the tools available to study brain activity in humans, arguably the most common method used to investigate attention is electroencephalography (EEG). Hundreds of studies have reported effects of attention on EEG recordings, both in frequency and time domains, some of which are highly robust; however, it is still not entirely clear what neural processes these differences represent.

Studies that leverage data in the frequency domain typically involve presenting temporally periodic stimuli and using Fourier analysis of the recorded EEG recordings to estimate power at the stimulus frequency. For example, steady-state visually evoked potentials (Friman et al., 2007; Norcia et al., 2015) and frequency tagging (Regan & Heron, 1969). This approach has several benefits, including being robust to noise (Srinivasan et al., 2006), and has been used to track the focus of attention (Toffanin et al., 2009). However, it necessarily forfeits the temporal dynamics of signal changes associated with attention, which can obscure the underlying mechanism.

EEG studies of attention in the time domain usually entail presenting stimuli to participants and analyzing common periods of neural activity around the presentations (i.e., epochs), which target the stimulus event-related potentials (ERP). The classical, univariate, analytical approach has identified ERP components that can be modulated by task demands associated with attention, e.g., the P3 (Polich & Bondurant, 1997; Sutton et al., 1965; Verleger, 1988). Univariate analysis of ERPs preserves the temporal dynamics of changes in the neural signals associated with attention, which can be used to infer underlying mechanisms. For example, by distinguishing feedforward from other processes, e.g., lateral and feedback (Zhang & Luck, 2009). However, it can be challenging to interpret how cognitive processes such as attention alter the neural representation of stimuli from univariate variations in sensor voltages.

More recently, several studies have used multivariate decoding of EEG recordings to study attention. Unlike univariate analyses, this method considers the pattern of signals (across the scalp and/or time) and can be used to measure the difference between the neural representation of two or more stimuli (Alilović et al., 2019; Bae & Luck, 2018). This method has been used to study both spatial and feature-based attention. For example, signals evoked by a range of visual stimuli are more reliably distinguished (i.e., accurately decoded) from ∼200 ms following stimulus presentation when they share a feature (e.g., colour/orientation) that is task relevant (Goddard et al., 2022; Grootswagers et al., 2021; Smout et al., 2019). Similarly, task-relevant visual stimuli can be more accurately decoded when their spatial location is cued prior to presentation (Foster et al., 2020, 2021). This increased decoding accuracy is not explained by stimulus expectancy nor engagement of working memory (Moerel et al., 2022), supporting the notion that it reflects the deployment of attention. The change associated with task relevance may reflect reduced neural variability of the stimulus representation (Arazi et al., 2019), for example, through an increase in the gain of attended features (Luck et al., 1997). However, increased decoding accuracy could also be produced by an additional cognitive process, which is prompted by the task relevant (target) stimulus feature and may not reflect feature-specific changes to the neural representation of the stimulus.

To test whether the influence of task relevance on EEG recordings reflects a change to the sensory representation of the attended stimulus, here we used a recently developed implementation of a multivariate analysis known as *inverted encoding* to decode EEG recordings of responses to coloured stimuli presented at different spatial locations (Brouwer & Heeger, 2009; Harrison et al., 2023; Rideaux et al., 2023). Specifically, we used this method to test whether spatial and feature-based attention change the neural representation of the same stimulus by making either the spatial location or colour of stimulus task relevant. For both spatial and feature-based tasks, we replicated the increased decoding accuracy that has been reported in previous studies (Foster et al., 2020, 2021; Goddard et al., 2022; Grootswagers et al., 2021; Smout et al., 2019). By comparing the effects associated with spatial and feature-based attention, we show that the change in the EEG recordings associated with task relevance is not feature specific. Indeed, our findings indicate that this phenomenon reflects an additional cognitive process, such as target recognition.

## METHOD

### Participants

Forty-two neurotypical human adults (mean±standard deviation age, 20.9±2.9 years; 10 males, 30 females, 2 non-binary) participated in the experiment. The data of one participant was omitted from analysis due to excessive eye movements. Sample size was informed by previous studies using similar neural decoding methods (Harrison et al., 2023; Rideaux et al., 2023). Observers were recruited from The University of Sydney and had normal or corrected-to-normal vision (assessed using a standard Snellen eye chart). All participants were naïve to the aims of the experiment and gave informed written consent. The experiment was approved by The University of Sydney Human Research Ethics Committee.

### Apparatus

The experiment was conducted in a dark, acoustically and electromagnetically shielded room. The stimuli were presented on a 24-inch ASUS VG248QE Gaming Monitor (ASUS, Taipei, Taiwan) with 1920 x 1080 resolution and a refresh rate of 144 Hz. The monitor was calibrated using a SpyderX Pro (Datacolor, Lawrenceville, NJ). Viewing distance was maintained at 70 cm using a forehead and chinrest, meaning the screen subtended 41.47° x 23.33° (each pixel 1.3’ x 1.3’). Stimuli were generated in MATLAB v2020a (The MathWorks, Inc., Matick, MA) using Psychophysics Toolbox (Brainard, 1997; Pelli, 1997) v3.0.18.13 (see http://psychtoolbox.org/). EEG was recorded on a 64 channel BrainVision ActiCap system (Brain Products GmbH).

### Stimuli, task, and procedure

The stimuli comprised coloured arcs (inner ring, 0.25°; outer ring, 4.25°) extending 45° polar angle on a mid-grey background. A centrally positioned black fixation dot (radius 0.25°) was presented to reduce eye movements. Trials consisted of 30 arc stimuli (location and colour randomly selected between 0-360°) presented for 0.2 s each, separated by a blank 0.1 s inter-stimulus-interval. The colour angle referenced values that were drawn from a circle in the CIE *L*\**a*\**b*\* colour space. This circle was centred in the colour space at (*L* = 70, *a* = 20, *b* = 38) with a radius of 60 (Zhang & Luck, 2008). Participants were then instructed to use the arrow keys to indicate the number of target stimuli that were presented during the preceding sequence. Task accuracy was calculated as the difference between the number of presented and reported targets, with stimuli within 10° of the target location/colour were considered targets. Given this criterion, the probability of target presentation was 20/360=.056. This was repeated 8 times per block. Participants performed 18 blocks of trials (∼60 min), receiving feedback on their detection accuracy at the end of each block. At the beginning of each block, target stimuli were identified as either those appearing at a defined location or colour, pseudo-randomly counterbalanced across the session. Target locations/colours were held constant for each participant, but pseudo-randomly counterbalanced across participants, to approximate a uniform distribution of target locations/colours between 0-360°.

### EEG

The EEG recordings were digitised at 1024 Hz sampling rate with a 24-bit A/D conversion. The 64 active scalp Ag/AgCl electrodes were arranged according to the international standard 10–20 system for electrode placement (Oostenveld & Praamstra, 2001) using a nylon head cap and with an online reference of FCz. Offline EEG pre-processing was performed using EEGLAB v2021.1 (Delorme & Makeig, 2004). The data were initially down-sampled to 512 Hz and subjected to a 0.1 Hz high-pass filter to remove slow baseline drifts and a 45.0 Hz low-pass filter to remove high-frequency noise/artifacts. Data were then re-referenced to the common average before being epoched into segments around each stimulus (-0.2 s to 1.5 s from the stimulus onset). The stimuli were presented rapidly ∼3 Hz, so epochs contained ERPs produced by multiple stimuli. However, stimuli were intentionally randomly sampled, such these additional signals could not provide any information that could be used to decode the current stimulus.

### Neural Decoding

To characterise sensory representations of the stimuli, we used an inverted modelling approach to reconstruct either the location or the colour of the arcs from the EEG recordings (Brouwer & Heeger, 2011). A theoretical (forward) model was nominated that described the measured activity in the EEG sensors given the location/colour of the presented arc. The forward model was then used to obtain the inverse model that described the transformation from EEG sensor activity to stimulus location/colour. For the main decoding analyses, the forward and inverse models were obtained using a ten-fold cross-validation approach in which 90% of the data were used to obtain the inverse model on which the remaining 10% were decoded. For the generalization analyses, the forward and inverse models were obtained using all the data from one condition and used to decode data from another condition.

Similar to previous work (Brouwer & Heeger, 2009), the forward model comprised five hypothetical channels, with evenly distributed idealized location/colour preferences between 0° and 360°. Each channel consisted of a half-wave rectified sinusoid raised to the fifth power. The channels were arranged such that a tuning curve of any location/colour preference could be expressed as a weighted sum of the five channels. The observed EEG activity for each presentation could be described by the following linear model:

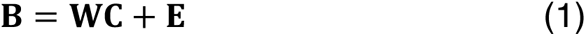

where **B** indicates the (*m* sensors × *n* presentations) EEG data, **W** is a weight matrix (*m* sensors × 5 channels) that describes the transformation from EEG activity to stimulus location/colour, **C** denotes the hypothesized channel activities (5 channels × *n* presentations), and **E** indicates the residual errors.

To compute the inverse model, we estimated the weights that, when applied to the data, would reconstruct the underlying channel activities with the least error. In line with previous magnetencephalography work (Kok et al., 2017; Mostert et al., 2015), when computing the inverse model, we deviated from the forward model proposed by (Brouwer & Heeger, 2009) by taking the noise covariance into account to optimize it for EEG data, given the high correlations between neighbouring sensors. We then estimated the weights that, when applied to the data, would reconstruct the underlying channel activities with the least error. Specifically, **B** and **C** were demeaned such that their average over presentations equalled zero for each sensor and channel, respectively. The inverse model was then estimated using either a subset selected through cross-fold validation) or all the data in one condition. The hypothetical responses of each of the five channels were calculated from the training data, resulting in the response row vector **c**_*train,i*_ of length *n*_*train*_ presentations for each channel *i*. The weights on the sensors **w***_i_* were then obtained through least squares estimation for each channel:

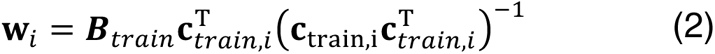

where **B**_*train*_ indicates the (*m* sensors × *n*_*train*_ presentations) training EEG data. Subsequently, the optimal spatial filter **v***_i_* to recover the activity of the *i*th channel was obtained as follows (Mostert et al., 2015):

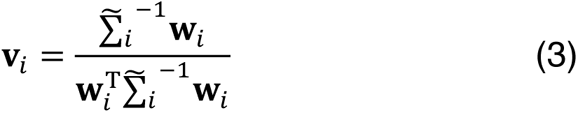

where 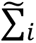 is the regularized covariance matrix for channel *i*. Incorporating the noise covariance in the filter estimation leads to the suppression of noise that arises from correlations between sensors. The noise covariance was estimated as follows:

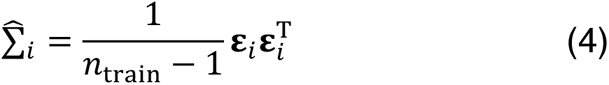

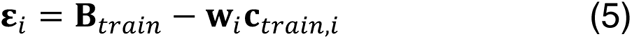

where *n*_train_ is the number of training presentations. For optimal noise suppression, we improved this estimation by means of regularization by shrinkage using the analytically determined optimal shrinkage parameter (Mostert et al., 2015), yielding the regularized covariance matrix 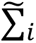.

For each presentation, we decoded location and colour by converting the channel responses to polar form:

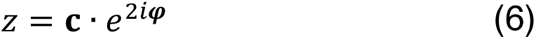

and calculating the estimated angle:

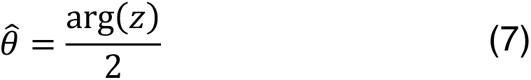

where **c** is a vector of channel responses and **φ** is the vector of angles at which the channels peak.

Stimulus location and colour properties were sampled from continuous distributions, but to reliably characterize these features across their dimension, we grouped presentations into 36 evenly spaced bins from 0-360°, relative to each participant’s attended location/colour. From the decoded locations/colours, we computed two estimates: *accuracy* and *precision*. Accuracy represented the similarity of the decoded location/colour to the presented location/colour (Kok et al., 2017), and was expressed by projecting the mean resultant (averaged across presentations within the same stimulus location/colour bin) of the difference between decoded and arc location/colour onto a vector with 0°:

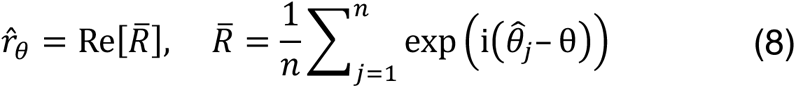

Precision was estimated by calculating the angular deviation (Zar, 1999) of the decoded locations/colours within each location/colour bin:

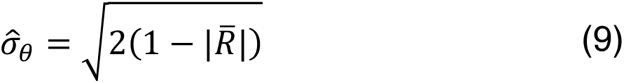

and normalized, such that values ranged from 0 to 1, where 0 indicates a uniform distribution of decoded location/colour across all location/colour (i.e., chance-level decoding) and 1 represents perfect consensus among decoded location/colour:

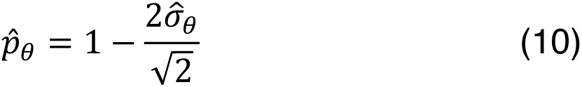

Prior to the main neural decoding analyses, we established the sensors that contained the most location/colour information by treating time as the decoding dimension and obtaining inverse models for each sensor, using ten-fold cross-validation. This analysis revealed that location/colour was primarily represented in posterior sensors (**Fig. 1a**); thus, for all subsequent analyses we only included signals from the parietal, parietal-occipital, and occipital sensors.

**Figure 1.**
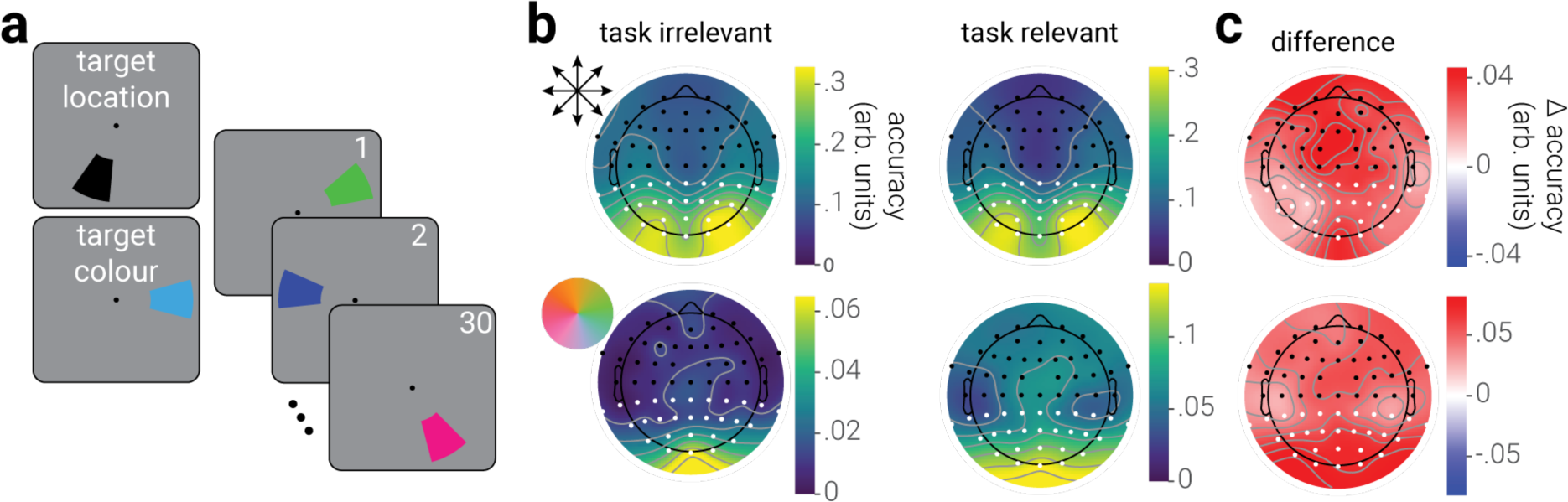
Experimental design and topographical distribution of stimulus information. **a**) Participants viewed blocks of 30 rapidly presented arcs (random polar angle and colour) and were tasked with reporting the number of target arcs, which were defined by either a particular location or colour. **b**) Decoding accuracy of the stimulus location (top row) and colour (bottom row) from the EEG recordings in blocks where the decoded stimulus property was either task irrelevant or relevant, across sensors. **c**) The difference between task irrelevant and relevant decoding accuracy for location and colour. White dots in (**b, c**) indicate the sensors from which information was used in all subsequent analyses.

### Eye tracking

Gaze direction recordings were epoched to the duration of stimulus presentations. To reduce the influence of eye movements on the results, stimulus presentations in which participants’ gaze direction exceeded 2° eccentricity from fixation were omitted from analyses (average±standard deviation omitted presentations, 4.2±8.0%). The data from one participant was omitted from analyses due to excessive eye movements, characterized as having exceeded the threshold for omission on >33% of presentations.

### Statistical analyses

Statistical analyses were performed in MATLAB v2020a and CircStat Toolbox v1.12.0.0 (Berens, 2009). For analyses of parameter estimates as a function of location/colour and time, a two-dimensional circular-linear cluster correction was applied to remove spurious significant differences. First, at each location/colour and time point, the effect size of accuracy/precision was calculated. Next, we calculated the summed value of these statistics (separately for positive and negative values) within contiguous featural-temporal clusters of significant values. We then simulated the null distribution of the maximum summed cluster values using permutation (*n*=1000) of the sign or condition labels, for single and paired *t*-test comparisons, respectively, from which we derived the 95% percentile threshold value. Clusters identified in the data with a summed effect-size value less than the threshold were considered spurious and removed. Prior to cluster analysis, we smoothed the parameter estimates along the feature dimension using a uniform kernel (size, 4).

### Data Availability

The EEG data generated in this study have been deposited in the following OSF database: https://osf.io/3k9cx/

## RESULTS

To investigate if attention shapes the neural representation of visual features, we recorded observers’ brain activity using EEG while they viewed blocks of rapidly presented coloured arcs. Observers were instructed to report the number of target arcs in each block, which were defined either by their spatial location or colour, as indicated at the beginning of the block (**Fig. 1a**). The location and colour of stimuli were selected at random. Thus, stimuli in target location blocks served to produce brain activity associated with spatial location being task relevant and colour being task irrelevant, and those in target colour blocks produced activity associated with colour being task relevant and spatial location being task irrelevant.

For all analyses of EEG recordings, we used inverted encoding to decode the visual properties (location or colour) of presented stimuli. From the trial-by-trial decoded properties, we derived summary parameter estimates of *accuracy* (the similarity between the decoded and presented stimulus property) and *precision* (the trial-by-trial reliability of the decoded property) as a function of time and property dimension.

We first computed the spatial configuration of the stimulus representation across the scalp by decoding location and colour information separately for each EEG sensor (**Fig. 1b**). We found that both stimulus properties were primarily represented in posterior sensors; however, location information was represented bilaterally while colour information was concentrated at the midline. For both properties, there was higher decoding accuracy for the corresponding task relevant condition across all sensors. The difference in accuracy between task relevant and irrelevant conditions was concentrated at posterior sensors (**Fig. 1c**); however, there was also a cluster of fontal electrodes that showed increased decoding accuracy of location in the task relevant condition, which could indicate the contribution of eye movements. To reduce the possible influence of eye movements, in all subsequent analyses, we omitted trials in which observers’ gaze deviated >2° from fixation and only analyzed EEG recordings from posterior sensors (**Fig 1b, c**, white dots).

We next computed decoding accuracy and precision as a function of time and distance from the target stimulus property. We found that the accuracy of spatial location increased sharply from ∼60 ms following stimulus onset and was relatively stable across time and robust to additional incoming information, i.e., subsequently presented stimuli, consistent with previous work (King & Wyart, 2021; Rideaux et al., 2023; **Fig. 2a**). However, task relevance increased the duration for which decoding accuracy was sustained, from ∼1000 ms to ∼1400 ms (**Fig. 2b**). Specifically, task relevance significantly increased the decoding accuracy of the target stimulus location from ∼200 - 1200 ms following stimulus presentation (**Fig. 2c**). We found a similar pattern of results for colour. However, the decoding accuracy of colour increased later (at ∼100 ms) than that for location and was only sustained for ∼500 ms when task irrelevant (**Fig. 2d, e**). Further, task relevance significantly increased accuracy for colour at both the target and opposite-to-target colour, producing a pattern of increased accuracy in the shape of a Mexican hat wavelet, centered on the target colour (**Fig. 2f**). The difference in accuracy between task irrelevant and relevant conditions was larger for colour. This may indicate that attention was more strongly engaged when detecting colour targets, for example, because it was more challenging than detecting targets based on location. Indeed, we found that (behavioural) detection accuracy for targets defined by location was significantly better than when defined by colour (*t*_40_=3.31, *p*=.002).

**Figure 2.**
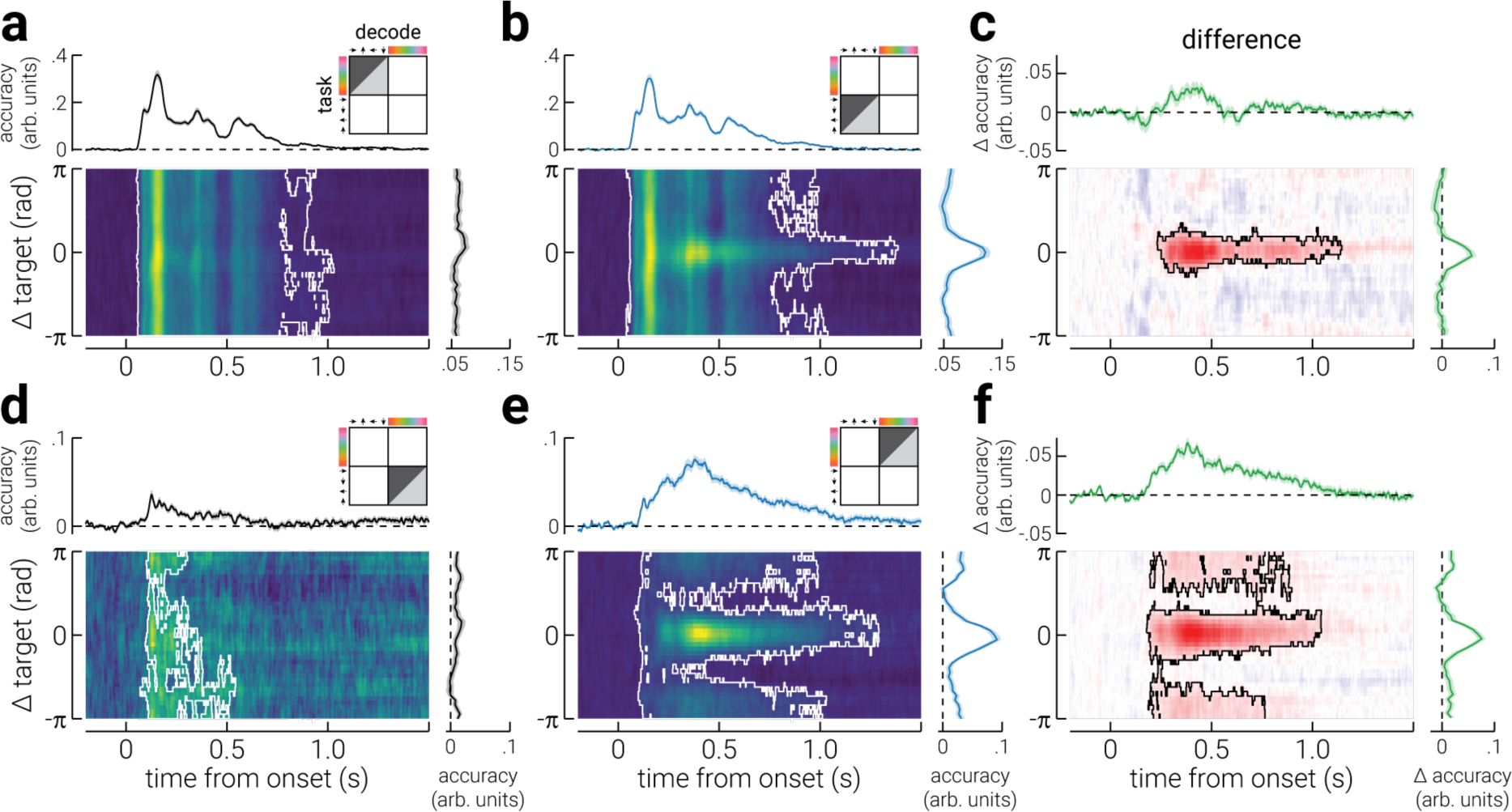
Accuracy of the neural representation of location and colour across time. **a**) A heatmap of the decoding accuracy for all spatial locations in the task irrelevant blocks (relative to participants’ target location) as a function of time (relative to stimulus onset); the white outline denotes cluster-corrected regions of statistical significance. The plots above and to the right of the heatmap show the temporal dynamics of accuracy (averaged across all locations) and the location-specific differences in accuracy (averaged across time), respectively. The illustration in the top-right corner indicates the task block/s data used to train (light grey triangle) and test (dark grey triangle) the decoder, and the decoded stimulus property. **b**) Same as (**a**), but for blocks where stimulus location was task relevant. **c**) The difference in decoding accuracy between (**b**) task irrelevant and (**a**) relevant conditions. **d-f**) The same as (**a-c**), but for colour. Note, the shaded regions in the plots positioned above and to the right of heatmaps indicate ±SEM. The temporally averaged accuracy plots do not include data prior to stimulus onset. The colour ranges used in the heatmaps (**a,b,d,e**) are normalized to the minimum and maximum values, to facilitate interpretation of relative differences; the adjacent plots should be used to gauge absolute values. The colour range used in (**c,f**) are normalized to ± absolute maximum value.

For decoding precision, we found a similar pattern of results as those for accuracy (**Fig. 3**). However, decoding precision in the task relevant conditions was only sustained until ∼1000 ms following stimulus onset (**Fig. 3b, e**), and the precision of colour did not significantly exceed chance in the task irrelevant condition (**Fig. 3d**).

**Figure 3.**
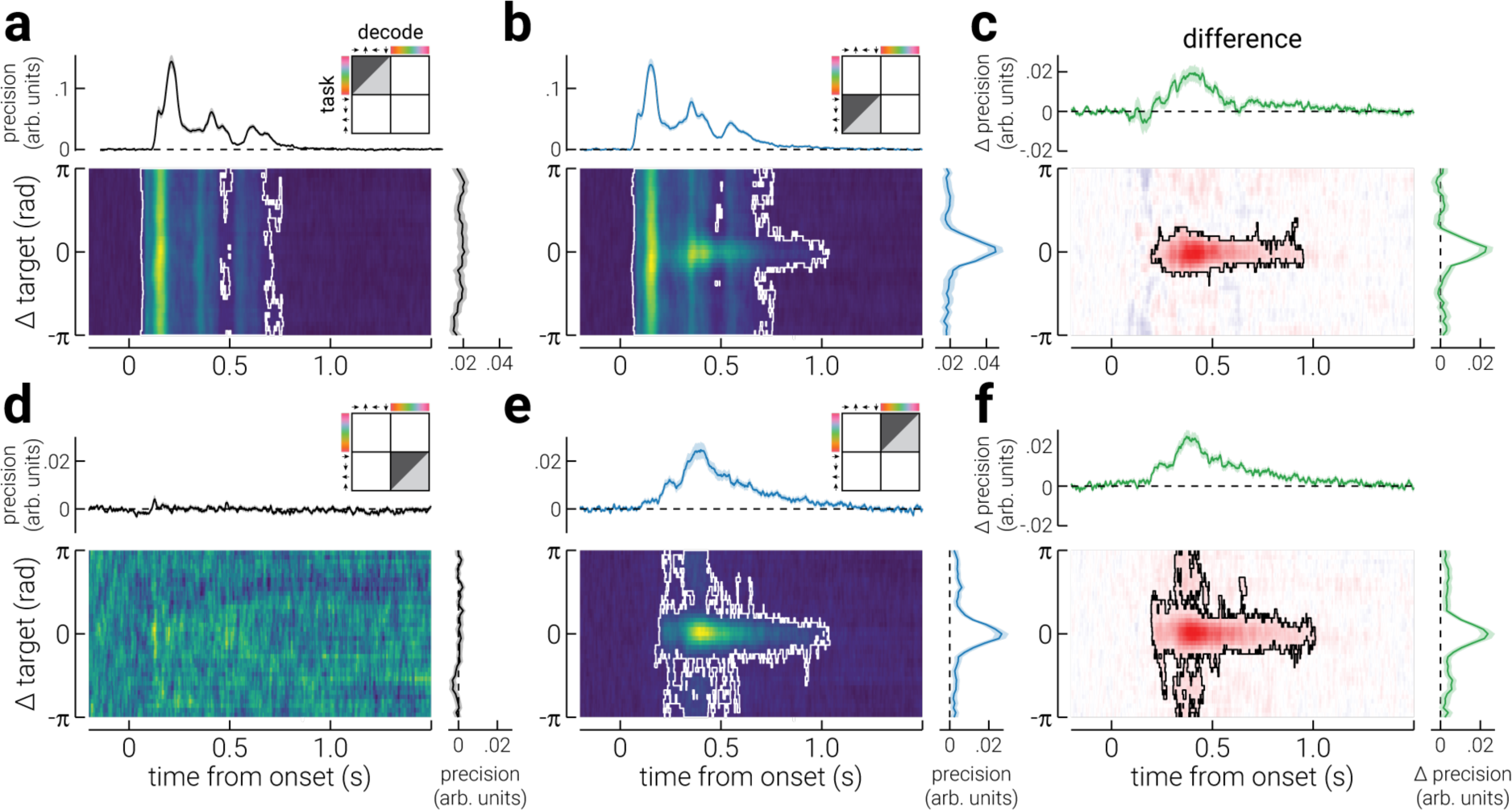
Precision of the neural representation of location and colour across time. **a**) A heatmap of the decoding precision for all spatial locations in the task irrelevant blocks (relative to participants’ target location) as a function of time (relative to stimulus onset); the white outline denotes cluster-corrected regions of statistical significance. The plots above and to the right of the heatmap show the temporal dynamics of precision (averaged across all locations) and the location-specific differences in precision (averaged across time), respectively. The illustration in the top-right corner indicates the task block/s data used to train (light grey triangle) and test (dark grey triangle) the decoder, and the decoded stimulus property. **b**) Same as (**a**), but for blocks where stimulus location was task relevant. **c**) The difference in decoding precision between (**b**) task irrelevant and (**a**) relevant conditions. **d-f**) The same as (**a-c**), but for colour. Note, the shaded regions in the plots positioned above and to the right of heatmaps indicate ±SEM. The temporally averaged precision plots do not include data prior to stimulus onset. The colour ranges used in the heatmaps (**a,b,d,e**) are normalized to the minimum and maximum values, to facilitate interpretation of relative differences; the adjacent plots should be used to gauge absolute values. The colour range used in (**c,f**) are normalized to ± absolute maximum value.

Consistent with previous work (Goddard et al., 2022; Grootswagers et al., 2021; Smout et al., 2019), we found that task relevance increases and sustains the decoding accuracy of visual properties (spatial location and colour) from EEG recordings from ∼200 ms following stimulus onset. We also found that decoding precision increased during this period, demonstrating that task relevance increases the test-retest reliability of the neural signal. The temporal dynamics of the effect appears similar for both spatial location and colour tasks, despite a considerable delay in the emergence of the neural representations of colour, relative to location.

The increase in decoding accuracy associated with task relevance has previously been attributed to improved encoding precision of the target feature (Ungerleider, 2000). If this were true, we would not expect the pattern of activity evoked by a target stimulus property (e.g., up/red) to qualitatively change when attended, but for it to be elicited more reliably, i.e., less test-retest variability (Arazi et al., 2019). To investigate this interpretation of the effect of task relevance, we computed the accuracy and precision of the neural signals in the task relevant conditions using inverted models trained on data from the task irrelevant conditions. For both location and colour, we found that task relevance did not significantly increase decoding accuracy (**Fig. 4a-d**). Rather, task relevance led to a small decrease in accuracy for spatial location (**Fig. 4b**). By contrast, for both location and colour, decoding precision was significantly increased by task relevance from ∼200 ms around the target stimulus property (**Fig. 4e-h**). The finding that task relevance increased the decoding precision, but not the accuracy, of spatial location and colour when using inverted models trained on the task irrelevant properties, shows that the pattern of neural activity associated with the task relevance stimulus properties became more reliable, but qualitatively different from that elicited by the task irrelevant properties.

**Figure 4.**
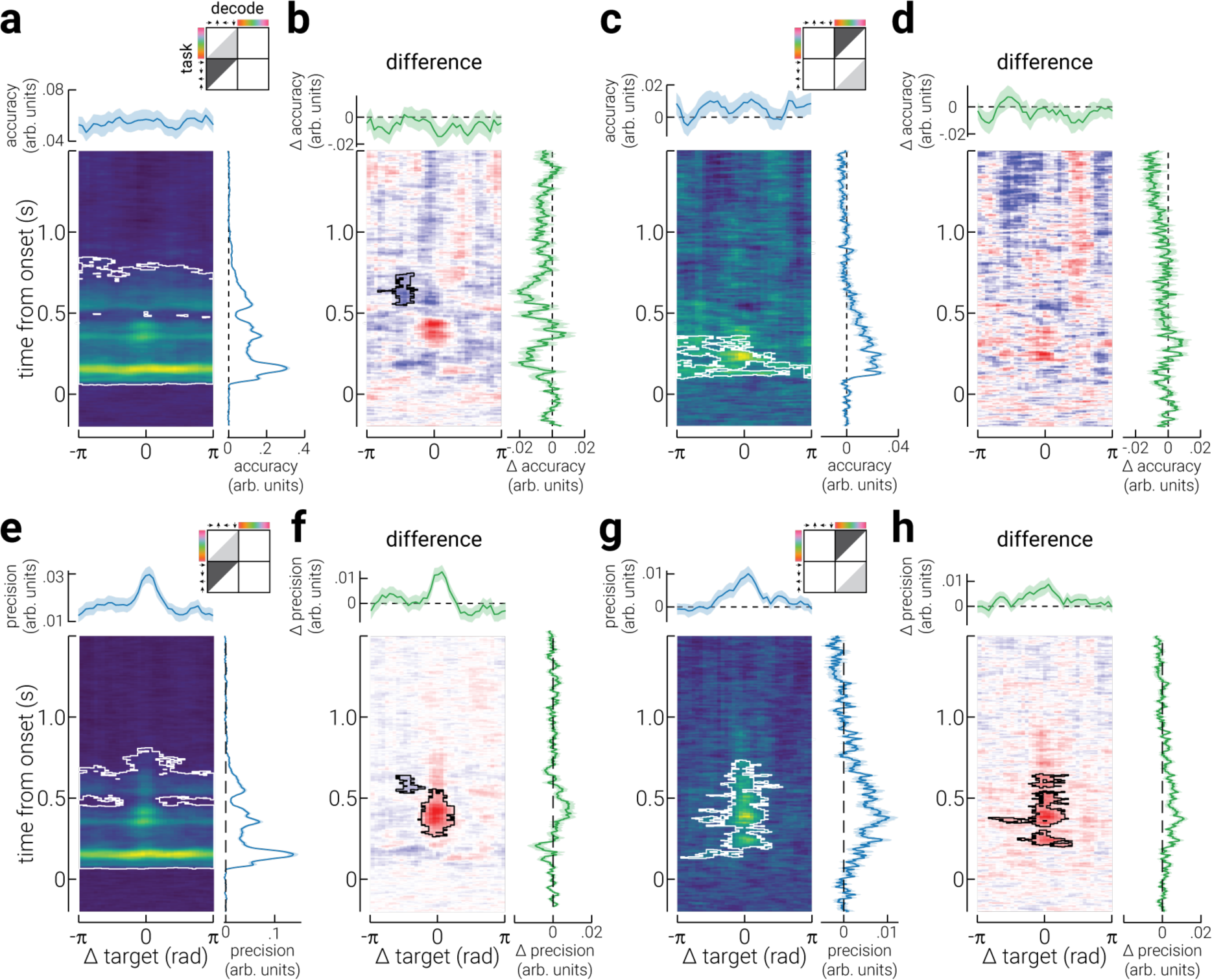
Generalizability between task irrelevant and relevant neural representations. **a**) A heatmap of the decoding accuracy for all spatial locations in the task relevant blocks as decoded by a model trained on data from the task irrelevant blocks. Accuracy is shown as a function of location (relative to participants’ target location) and time (relative to stimulus onset); the white outline denotes cluster-corrected regions of statistical significance. The plots above and to the right of the heatmap show the location-specific differences in accuracy (averaged across time) and the temporal dynamics of accuracy (averaged across all locations), respectively. The illustration in the top-right corner indicates the task block/s data used to train (light grey triangle) and test (dark grey triangle) the decoder, and the decoded stimulus property. **b**) Same as (**a**), but for the difference between accuracy in task irrelevant and relevant blocks, decoded with a model trained on signals from task irrelevant blocks. **c, d**) Same as (**a, b**), but for colour. **e-h**) Same as (**a-d**), but for decoding precision. Note, the shaded regions in the plots positioned above and to the right of heatmaps indicate ±SEM. The temporally averaged accuracy and precision plots do not include data prior to stimulus onset. The colour ranges used in the heatmaps (**a,c,e,g**) are normalized to the minimum and maximum values, to facilitate interpretation of relative differences; the adjacent plots should be used to gauge absolute values. The colour range used in (**b,d,f,h**) are normalized to ± absolute maximum value.

The qualitative change in the pattern of neural activity associated with a stimulus property being task relevant could reflect an additional cognitive process, e.g., recognition of a target stimulus. If this were true, we would expect that a similar pattern of activity would be produced by both tasks. To test this, we normalized all data to participants’ target property, i.e., the target colour and location were positioned 0°, and decoded each stimulus property using an inverted model trained to decode the other property. On each presentation, the stimulus location and colour were selected randomly and therefor orthogonal. Thus, as expected, for the task irrelevant properties we found that decoding accuracy did not significantly exceed chance (**Fig. 5a, d**). By contrast, the accuracy of task relevant properties significantly exceeded chance from ∼200 ms following stimulus onset, particularly at the target property (**Fig. 5b, e**). These results show that the pattern of activity elicited by task relevant demands, which emerges at ∼200 ms following stimulus onset, is not feature specific.

**Figure 5.**
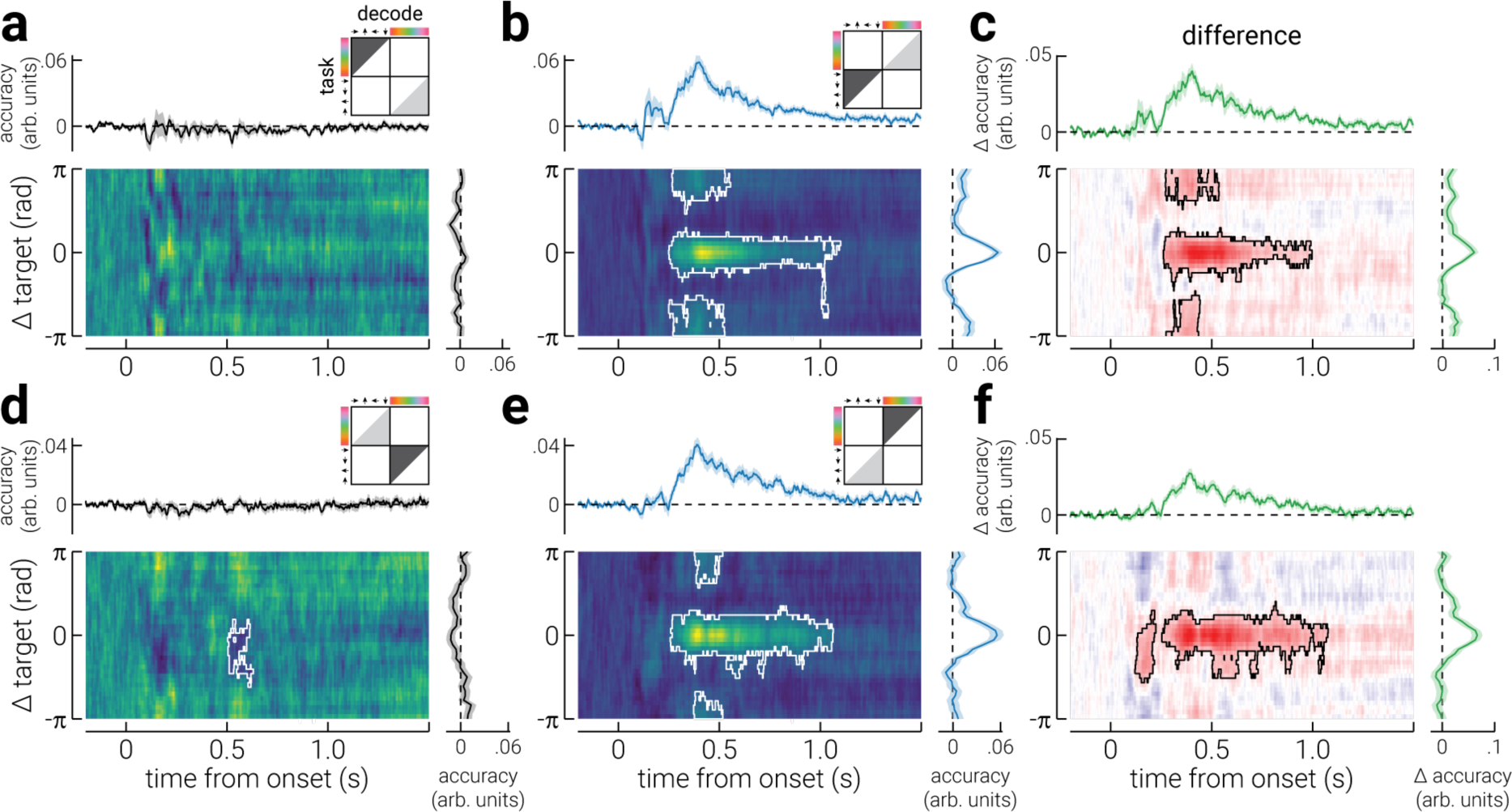
Generalizability between stimulus properties. **a**) A heatmap of the decoding accuracy for all spatial locations in the task irrelevant blocks as decoded by model trained to decode colour in the unattended blocks. Accuracy is shown as a function of location (relative to participants’ target location) and time (relative to stimulus onset); the white outline denotes cluster-corrected regions of statistical significance. The plots above and to the right of the heatmap show the temporal dynamics of accuracy (averaged across all locations) and the location-specific differences in accuracy (averaged across time), respectively. The illustration in the top-right corner indicates the task block/s data used to train (light grey triangle) and test (dark grey triangle) the decoder, and the decoded stimulus property. **b**) Same as (**a**), but for blocks where stimulus location was task relevant. **c**) The difference in decoding precision between (**b**) task irrelevant and (**a**) relevant conditions. **d-f**) The same as (**a-c**), but for colour. Note, the shaded regions in the plots positioned above and to the right of heatmaps indicate ±SEM. The temporally averaged accuracy plots do not include data prior to stimulus onset. The colour ranges used in the heatmaps (**a,b,d,e**) are normalized to the minimum and maximum values, to facilitate interpretation of relative differences; the adjacent plots should be used to gauge absolute values. The colour range used in (**c,f**) are normalized to ± absolute maximum value.

## DISCUSSION

Attention supports efficient perception by increasing the neural signals of target locations/features while supressing the those of distractors. The influence of task demands that engage attention can be reliably detected within EEG recordings. Decades of univariate ERP analyses have provided foundational understanding of attention in the human brain, but many aspects of this phenomenon remain unknown. Multivariate analyses may provide a means of addressing these questions. EEG work using multivariate linear decoding has reliably demonstrated increased decoding accuracy for task relevant features from ∼200 ms following stimulus onset (Goddard et al., 2022; Grootswagers et al., 2021; Smout et al., 2019), which is not explained by working memory load or expectation (Moerel et al., 2022). While some work indicates that this effect is not linked to feedforward processing (Alilović et al., 2019), other work has used this effect to distinguish between the influence of spatial and feature-based attention on stimulus representation (Goddard et al., 2022). Neurophysiological work in macaque has shown that attention alters the representation of stimuli in sensory cortices (Brefczynski & DeYoe, 1999; Gandhi et al., 1999; Luck et al., 1997; Motter, 1993; Treue & Maunsell, 1996). However, it is unclear whether the increased decoding accuracy from ∼200 ms represents a change in the stimulus representation or an additional cognitive process. To understand what the change in multivariate information that is associated with task demands reflects, here we used inverted encoding to characterize how spatial and feature-based tasks shape the representation of space and colour.

We found that task relevance increased decoding accuracy from ∼200 ms following stimulus onset, for both spatial location and colour. We further found that the difference in decoding accuracy formed a Gaussian profile when assessed as a function of the distance from the target stimulus property, such that it was maximally improved at the target location/colour. We found a similar pattern of results for decoding precision, which provides a direct index of the test-retest reliability of the neural response. These findings seemed to support the interpretation that the increased decoding accuracy for task relevant properties reflects more stable stimulus representations. However, this interpretation seems unlikely for two reasons. First, the difference in decoding accuracy and precision between task relevant and task irrelevant conditions appears ∼150 ms after location and colour can be reliably decoded, indicating that the resolution of the sensory representations are the same between conditions during the feedforward sweep of visual processing. Second, the results from the generalization analysis, shown in **Figure 5**, show that there is a neural signal that begins at the same time as these changes (∼200 ms), and is invariant to the visual feature. These findings provide compelling evidence that task-related change in ERPs do not reflect a change in the stimulus representation. Rather, the increased decoding accuracy of task relevant features from EEG recordings would appear to reflect a change in the neural representation that is stimulus invariant. A change in the neural representation of this kind likely represents an additional cognitive process, such as template matching (Desimone & Duncan, 1995; Myers et al., 2015).

The biased competition framework posits that attention tonically pre-activates visual neurons with receptive fields tuned to template-matching stimuli (Reynolds & Chelazzi, 2004). Increasing the baseline activity of stimulus-specific representations is thought to facilitate target selection and reduce distractor competition for downstream processing resources (Bundesen et al., 2005; Maunsell & Treue, 2006). Myers et al. (2015) tested the MEG and EEG signatures of template matching by decoding top-down mnemonic template and bottom-up stimulus features during a detection task. While visual features (orientation) could be decoded as early as 50 ms following stimulus presentation, the difference between the target feature and stimulus feature could only be decoded from around ∼200 ms. The timing of their decoding results is consistent with those observed here and in other EEG decoding studies and seems to be very robust (Goddard et al., 2022; Grootswagers et al., 2021; Smout et al., 2019). However, the lack of any task-relevant modulation during the initial feedforward sweep seems to refute the claim that attention pre-activates early cortical neurons. While, the current results are most consistent with participants comparing incoming sensory information with a mnemonic template of the target feature, maintaining this template does not appear to alter the sensory representation. A match between the template and the stimulus produced an additional neural response that was the same regardless of whether the template and stimulus were defined by spatial location or colour.

Moerel et al. (2022) also found that orientation decoding accuracy at ∼200 ms after onset for colours bars was better when one of the bars was task relevant (e.g., horizontal blue). The researchers defined stimuli as either blue or orange and one of four possible orientations. To mitigate against the influence of target detection on the decoding results, they omitted the target orientation from the classification analysis. Despite the target orientation not being included in the decoding analysis, they still observed better decoding of orientation between stimuli with the target colour, compared to the non-target colour. How do we resolve their results with ours, which seem to suggest that improved decoding accuracy reflects target detection? Target detection for continuous features, e.g., orientation or spatial location, likely operates in a continuous manner. That is, when a stimulus is presented, there will be varying levels of correlation between the mnemonic template of the target and the stimulus. This is presumably true even when the feature is presented in a discretised manner, as in Moerel et al. (2022). Indeed, while we found decoding accuracy for both colour and spatial location peaked at the target colour/location, the improvement is also seen (to a lesser extent) for colours/locations around the target colour/location. We even found better decoding in the opposite-to-target colour/location (most pronounced for colour, but visible for spatial location as well), which may reflect a confident non-target recognition. Indeed, if we only consider the decoding accuracy for non-target colours/locations, like Moerel et al. (2022), we also find higher average decoding accuracy in the task-relevant condition compared to the task-irrelevant condition. Critically, however, we do not see transfer from unattended to attended features, indicating that the boost does not represent increased fidelity of the sensory representation. Thus, our results indicate that removing the influence of target recognition processes from neural activity in these tasks may be more challenging than removing responses to target stimuli.

While our findings may be inconsistent with some interpretations of recent studies using multivariate analyses (Goddard et al., 2022; Smout et al., 2019), they are consistent with results from univariate ERP work. In particular, the timing of the increased decoding (∼200 ms following stimulus onset) aligns with the onset of the P3 component (Polich & Bondurant, 1997; Sutton et al., 1965; Verleger, 1988). This component is reliably modulated by attention (Pfefferbaum et al., 1985), however, its topographical representation is relatively invariant to input modality (Ji et al., 1999). The P3 is also modulated by stimulus probability (Duncan-Johnson & Donchin, 1977). While we used a uniform distribution of stimuli, it is possible that observers grouped stimuli into target and non-target categories. Thus, while the probability of stimuli was equal, target stimuli were less frequent than non-target stimuli, which may have contributed to modulation of the P3 component. Given initial target recognition is necessary for this to have occurred (i.e., only by recognising that stimuli were or were not targets could their probability seem non-uniform), the increased decoding observed from ∼200 ms following stimulus onset is most parsimoniously explained by target recognition (i.e., template matching), but may also reflect some information about stimulus probability. Critically, however, the cognitive process associated with the changes in neural activity is stimulus invariant. While this signal is not suitable for studying the influence of attention on stimulus representation, further investigation of the process would provide further understanding of the cascade of cognitive processes associated with target detection.

One limitation of the current study is that we did not have a condition in which attention was directed away from the decoded stimuli, for example, by using a fixation task. Foster et al. (2021) found that decoding accuracy was higher for stimuli that were the focus of spatial attention, compared to a those where attention directed away. While they observed improved decoding accuracy in the earliest stages of sensory processing (∼80 ms), it is unclear to what extent the lingering activity from previous, correlated, stimuli confounded this result. Here we avoided this confound by presenting stimuli randomly, such that preceding and subsequent stimuli did not produce activity that could improve the decoding accuracy of the current stimulus. While we did not include a condition in which attention was directed away from all stimuli, in the spatial target condition, participants attention was directed away from stimuli outside the target location. Thus, if spatial attention altered the sensory representation of the attended stimuli, we would expect to see improved decoding accuracy for the target location, relative to other locations, during the feedforward sweep of the visual cascade, i.e., ∼50-100 ms, yet we only observed increased accuracy later (∼200 ms). Additionally, while we found no difference between spatial and feature-based attention, this may have been due to the visual properties we selected. Future work could test other visual properties, such as orientation.

Previous EEG work has found evidence that early ERP components are modulated by task relevance (Zhang & Luck, 2009), suggesting that changes to feedforward stimulus-evoked activity can be detected with EEG. By contrast, recent work applying multivariate analyses to EEG recordings found no evidence that spatial task relevance changed decoding accuracy during feedforward processing (Alilović et al., 2019). Similarly, here we found no evidence that task relevance (spatial or feature-based) altered feedforward processing. While our findings urge caution when interpreting multivariate decoding results, if future work can develop experimental paradigms that can produce reliable attention-related changes in activity evoked during feedforward processing, these data-driven analytic techniques combined with forward modelling may provide key insights into how spatial and feature-based attention influence sensory processing in humans.

